# Aging-Associated *Nox4*-Mediated Mitochondrial ROS and DNA Damage Promote Vascular Cell Reprogramming and Aortic Remodeling in Abdominal Aneurysms

**DOI:** 10.1101/2025.07.09.664017

**Authors:** Aleksandr E. Vendrov, Jamille Chamon, Julia Levin, Takayuki Hayami, Chaitanya Madamanchi, Morgan Salmon, Nageswara R. Madamanchi

## Abstract

**Background:** Aging and male sex are major risk factors for abdominal aortic aneurysm (AAA), a disease characterized by vascular cell phenotypic switching and aortic wall remodeling. Mitochondrial oxidative stress (mtOS) has been implicated in these changes. We previously demonstrated that NOX4 expression and activity increase with age in cardiovascular cells, promoting mtOS and vascular dysfunction. This study investigates whether NOX4-driven mtOS and DNA damage promote AAA development through vascular cell reprogramming.

**Methods:** We used mitochondria-targeted *Nox4*-overexpressing (*Nox4*TG) mice with an *Apoe^-/-^* background to model Angiotensin II (Ang II)-induced AAA. AAA incidence, aortic morphology, reactive oxygen species (ROS) levels, DNA damage markers, and wall remodeling parameters were assessed in *Apoe^-/-^*, *Apoe^-/-^*/*Nox4*TG, and *Apoe^-/-^/Nox4^-/-^* mice. Vascular cell populations were analyzed by spectral flow cytometry and gene expression profiling. In vitro, Ang II-treated SMCs from wild-type, *Nox4*TG, and *Nox4^-/-^* mice were evaluated for mtROS, DNA damage, and activation of inflammatory pathways.

**Results:** *Apoe^-/-^/Nox4TG* mice exhibited the highest AAA incidence, aortic dilation, ROS levels, DNA damage, and inflammation, while *Apoe^-/-^/Nox4^-/-^* mice were most protected. Macrophage-like SMCs increased, while contractile SMCs decreased in Nox4TG aortas. Ang II-treated *Nox4*TG SMCs showed elevated mtROS, DNA damage, and cGAS-STING activation. Flow cytometry analysis confirmed the presence of aneurysmal SMC with reduced ACTA2, MYH11, TAGLN and increased CD68, CD11b, LGALS3 expression.

**Conclusions:** NOX4-dependent mitochondrial DNA damage and activation of DNA-sensing pathways promote SMC phenotypic switching, inflammation, and aortic wall remodeling in AAA. Targeting NOX4 and enhancing mitochondrial function may offer therapeutic strategies for AAA prevention.

**CLINICAL PERSPECTIVE:** *What Is New?:*

- This study identifies mitochondrial NOX4-derived ROS and oxidative mitochondrial DNA damage as early and causal events in AAA pathogenesis, promoting pro-inflammatory reprogramming of vascular smooth muscle cells toward a macrophage-like phenotype.
- Activation of cytosolic DNA sensing (cGAS-STING) and DNase II pathways in response to mitochondrial damage links redox imbalance to innate immune activation and progressive aortic wall remodeling in AAA.

*What Are the Clinical Implications?:*

- Targeting mitochondrial-specific oxidative stress or downstream DNA-sensing pathways offers a novel therapeutic strategy to halt AAA progression in patients with small, asymptomatic aneurysms.
- These findings provide a mechanistic basis for developing mitochondria-targeted antioxidants or NOX4 inhibitors as potential pharmacologic interventions to delay or prevent AAA rupture, which could reduce the need for surgical repairs.

## INTRODUCTION

A ruptured abdominal aortic aneurysm (AAA) is the 13th most common cause of death in the United States. About 1% of all deaths in the West are caused by ruptured abdominal aortic aneurysms.^1^ Older age is a significant risk factor for the development of aortic aneurysms. Other risk factors include a family history of abdominal aortic aneurysm, coronary artery disease, hypertension, peripheral artery disease, and previous myocardial infarction.^2^ Men are 4-6 times likely than women to develop AA.^3^

For aneurysms larger than 5.5 cm, the only treatment options available are open surgical or endovascular repair.^4^ Current strategies for managing aneurysms smaller than 5.5 cm are inadequate, often leading to unnoticed growth and potentially fatal ruptures.^5^

The pathophysiology of AAA is complex and involves the degradation of the elastic media of the aortic wall, which causes aortic dilation and rupture. Although the exact molecular mechanisms underlying AAA are unclear, oxidative stress has been shown to play a causal role in experimental AAA and other types of aneurysms.^6,7^ A growing body of evidence suggests that mitochondrial oxidative stress (mtOS) plays a role in the development of aging and age-related diseases.^8,9^ The decrease in mitochondrial respiration, associated with mitochondrial dysfunction, can lead to normal arterial aging and age-related cardiovascular diseases, such as atherosclerosis and aneurysms.^10,11^ In patients with AAA, an increase in mitochondrial and endoplasmic reticulum stress, coupled with a reduction in mitochondrial biogenesis, was observed within the vascular walls.^12^

NADPH oxidases, specifically NOX1, NOX2, NOX4, and NOX5, are the primary NOX isoforms in the vascular tissue. However, NOX5 is absent in rodents.^13,14^ Both NOX1 and NOX2 are in the plasma membrane and require cytosolic subunits and p22phox for activation. In contrast, NOX4 is constitutively active and is found in intracellular compartments such as the endoplasmic reticulum and mitochondria.^15,16^ The activity of NOX4 is mainly regulated at the transcriptional level, and it is upregulated in response to various physiological and pathological stimuli.^17^ NOX4 primarily produces H₂O₂, a diffusible ROS that plays a crucial role in modulating redox-sensitive signaling pathways associated with vascular homeostasis and remodeling.^18^ A genome-wide association analysis has identified NOX4 as a likely causal gene in the development of AAA, suggesting it may play a role in the dysregulation of the extracellular matrix.^19^ However, direct functional evidence, such as experimental manipulation of NOX4 in cellular or animal models, is still necessary to confirm its role in the pathogenesis of AAA.

We previously demonstrated that NOX4 expression increases with age in cardiovascular tissues, promoting mitochondrial oxidative stress (mtOS) and contributing to vascular aging and disease.^16, 20–26^ However, the role of NOX4-induced mtOS and mitochondrial DNA (mtDNA) damage in inflammatory reprogramming, SMC phenotypic switching, and vascular immune responses in AAA pathogenesis has not yet been explored. To address this gap, we used *Nox4* transgenic (*Nox4*TG) and knockout (*Nox4*^-/-^) mice, crossed with *Apoe*^-/-^ mice, in a well-established model of Angiotensin II (Ang II)-induced AAA to dissect the role of NOX4-mediated mtOS in the remodeling of the aortic wall and the progression of the disease. **Notably, middle-aged *Nox4*TG mice exhibit cardiovascular aging features such as aortic stiffness,** ^21^ **increased susceptibilities to ventricular tachyarrhythmia,**^24^ **and diastolic dysfunction.**^23^

Our findings demonstrate that an increase in mitochondrial NOX4 expression in a hyperlipidemic background significantly increased Ang II-induced AAA. This increase is linked to a phenotypic change that drives the SMCs toward a pro-inflammatory, macrophage-like state, which is characterized by heightened cytokine production and the expression of immune-related genes. These findings indicate a previously unrecognized role for mitochondrial NOX4 in enhancing vascular inflammation by reprogramming SMCs.

## MATERIALS AND METHODS

### Human Samples

Deidentified human samples were not classified as human subjects research and were approved by the University of Michigan Institutional Review Board (IRB). Human tissue was collected following the guidelines set by the Human Subjects Review Committee at the University of Virginia (IRB protocol 12178). Preoperative consent was obtained from all patients. AAA tissue was resected from male patients during open surgical AAA repair, while abdominal aortic tissue from transplant-donor patients was collected to serve as controls.

### Animals

All animal procedures complied with the protocols approved by the University of Michigan Institutional Animal Care and Use Committee, following NIH OLAW policy and Guide for the Care and Use of Laboratory Animals. Male wild-type C57BL/6J mice, *Nox4*TG (C57BL/6J-Tg(CAG-COX8A/Nox4)618Mrng/J),^21^ *Nox4*^-/-^ (B6.129-*Nox4^tm1Kkr^*/J),^25^ and *Apoe*^-/-^ (B6.129P2-*Apoe^tm1Unc^*/J) mice were purchased from the Jackson Laboratory (Bar Harbor, ME). **In accordance with the recommendations of the AHA ATVB Council,**^27^ **only male mice were utilized in all experiments.** The mice were crossed and bred in-house, and littermate mice were used in all experiments. Mice were housed in ventilated cages at 22°C, with a 12-hour light/dark cycle, and had free access to food and water.

### AAA Model

Six-month-old male mice, including *Apoe*^-/-^, *Nox4*TG/*Apoe^-/-^,* and *Nox4^-/-^/Apoe*^-/-^, were surgically implanted with subcutaneous micro-osmotic pumps (Alzet 1002, Durect; Cupertino, CA) for continuous infusion of Angiotensin II (Ang II) at a dosage of 1000 ng/kg/min for 28 days. The morphology and size of the descending aorta were assessed using Doppler ultrasound before and 28 days after the administration of Ang II, employing the Vivo 2100 Imaging System (FUJIFILM VisualSonics Inc., Bothell, WA). After the study, the mice were euthanized with an overdose of inhaled isoflurane. The circulatory system was then perfused with phosphate-buffered saline, and the entire aortas were dissected, grossly examined, and either embedded in O.C.T. compound (Sakura Finetek, Torrance, CA), snap-frozen in liquid nitrogen, or used for cell suspension preparation.

### Cell Culture

Vascular smooth muscle cells were isolated from the aortas of wild-type, *Nox4*TG, and *Nox4*^-/-^ mice, following previously established methods. ^16^ The cell cultures were maintained in Dulbecco’s Modified Eagle’s Medium (DMEM), supplemented with 10% fetal bovine serum and antibiotic/antimycotic solution (Thermo Fisher, Waltham, MA), in a 5% CO_2_ incubator at 37°C. Cells were supplemented with serum-free DMEM for the experiments and treated with either vehicle or 100 µM Ang II.

### Histology and Immunostaining

Transverse 10 µm frozen sections of the abdominal aorta were stained using a Verhoeff-van Gieson elastic stain kit, following the manufacturer’s recommendations (Abcam; Waltham, MA). Two blinded observers scored elastin degradation in the aortic wall on the four-point scale (1 – mild disruption; 2 – moderate disruption; 3 – severe disruption; 4 - greater than 75% degradation). Picrosirius red staining was performed as previously reported using Picrosirius Red Stain Kit (Abcam).^26^

Immunofluorescence staining of cryosections from the abdominal aorta was performed as previously described.^26^ In brief, the cryosections were fixed in acetone and permeabilized with 0.1% Triton X-100. The immunostaining was conducted using the following antibodies: NOX4, 8-OHdG, TOMM20-AlexaFluor488, and IL1β (Abcam); FGA and DNASE2 (Thermo Fisher); ATP5G-AlexaFluor488, CD68-AlexaFluor488, CD68-Cy3, and MYH11-AlexaFluor488 (Bioss, Woburn, MA); ACTA2-FITC and ACTA2- Cy3 (from Sigma-Aldrich, St. Louis, MO); CD11b (Itgam) (Abnova, Walnut, CA); and CGAS, STING, and IL6 (Cell Signaling Technology, Danvers, MA). When appropriate, goat anti-rabbit secondary antibodies conjugated to AlexaFluor488 or AlexaFluor594 were used. The sections were mounted with ProLong Gold reagent containing DAPI (Thermo Fisher). Fluorescence images were captured using an Echo Revolve R4 microscope (Echo, San Diego, CA) and analyzed with NIH ImageJ 1.54f (Bethesda, MD).

### Western Blot and ELISA Analysis

Western blotting was performed as previously described.^28^ Briefly, total protein lysates were prepared from human or mouse aorta tissue using T-PER Tissue Protein Extraction Reagent or from mouse cells using M-PER Mammalian Protein Extraction Reagent supplemented with Halt Protease Inhibitor Cocktail (Thermo Fisher). Protein concentration was measured using Pierce BCA Protein Assay Kit (Thermo Fisher). The lysates were resolved on 10% SDS-PAGE and transferred to a nitrocellulose membrane (GE Healthcare, Chicago, IL). The primary antibodies used were NOX4 (Abcam), DNASE2 (Thermo Fisher), CGAS, STING, phosphor-STING (Cell Signaling), ACTB, and TUBB (Sigma). The secondary antibodies used were anti-mouse or anti-rabbit conjugated antibodies IRDye™ 800CW or IRDye™ 680CW (Li-Cor, Lincoln, NE). The specific band fluorescence signal was measured and analyzed using the Odyssey CLx imaging system (Li-Cor).

Lysed mouse VSMCs were tested for 8-OHdG expression using OxiSelect™ Oxidative DNA Damage ELISA Kit (Thermo Fisher) and for 2’3’-cGAMP using 2’3’- cGAMP EIA Kit (Cayman Chemical, Ann Arbor, MI), following the manufacturer’s protocols.

### ROS Detection

ROS levels were measured immediately after sample collection, following previously established methods. ^26^ Frozen human and mouse abdominal aorta sections were treated with 10 μM DHE or 5 μM MitoSOX Red (Thermo Fisher) for 15 minutes. Mouse VSMCs were also incubated with 5 μM MitoSOX Red for 15 minutes. Fluorescence images were captured using an Echo Revolve R4 microscope and analyzed with NIH ImageJ software. Images were analyzed with NIH ImageJ. To calculate the final DHE and MitoSOX Red fluorescence values, background fluorescence from PEG-SOD- treated controls was subtracted from the readings.

### Mitochondrial Function Analysis

The mitochondrial respiration of VSMCs was assessed using a Seahorse XFe96 analyzer (Agilent Technologies, Santa Clara, CA), following previously established protocols. ^21,26^ In brief, 50000 cells per well were plated on Seahorse XFe96 cell culture microplates and treated with either a vehicle or 100 µM Ang II for 24 hours. The oxygen consumption rate (OCR) was measured using the Seahorse XFp Cell Mito Stress Test Kit (Agilent). After the measurement, protein concentrations for each well were determined immediately using the Pierce BCA Protein Assay Kit (Thermo Fisher), and the OCR levels were normalized to the protein amount per well. Mitochondrial function parameters were calculated as previously described.^16^

### Spectral Flow Cytometry

The single-cell suspension from the mouse abdominal aorta was analyzed as previously described.^26,29,30^ A multi-color flow cytometry antibody panel was created using the FluoroFinder Panel Builder (FluoroFinder, Broomfield, CO). To prepare the samples, abdominal aorta segments were finely minced and dissociated in Hanks’ balanced salt solution (HBSS) containing 400 U/mL collagenase type I, 120 U/mL collagenase type XI, 60 U/mL hyaluronidase, and 60 U/mL DNAse I (Sigma) at 37°C. The resulting cell suspension was filtered through a 70 µm cell strainer (Corning, Corning, NY) and washed with FACS buffer (HBSS supplemented with 1% BSA and 1 mM EDTA). Samples were then resuspended in 200 µL FACS buffer containing the cell surface marker antibody mix, which included CD45-PE Fire 810, CD38-PE Fire 700, CD68-Brilliant Violet 785 (BioLegend, San Diego, CA), and CD11b-NovaFLuor Blur 610/30S (Thermo Fisher). After incubating the samples for 60 min at 4°C, they were washed and permeabilized in Foxp3/Transcription Factor Staining Buffer Set (Thermo Fisher) following the manufacturer’s instructions.

Samples were incubated in 200 µL FACS buffer with a mix of intracellular marker antibodies: (MYH11-FITC (Biorbyt, Durham, NC), CNN1-AlexaFluor 555 (Bioss), ACTA2-DyLight 405, TAGLN-PerCP (Novus Biologicals, Centennial, CO), TNFα-Brilliant Violet 510 (BioLegend), IL1b-APC-Cy7 (LifeSpan Bio, Newark, CA), and IL6-PerCP-eFluor 710 (Thermo Fisher). After a 60 min incubation, the samples were washed and resuspended in 300 µL FACS buffer. Flow cytometry analysis was performed using the Aurora Spectral Analyzer (Cytek, Freemont, CA) according to the manufacturer’s protocol. The negative control cell sample was left unstained. UltraComp Beads (Thermo Fisher), bound with corresponding antibodies, were used as single stain controls. The unmixing matrix with autofluorescence extraction was calculated using SpectroFlo (Cytek). All samples were analyzed using live unmixing.

Unmixed FCS files were analyzed using FlowJo 10 (BD Biosciences, Franklin Lakes, NJ) or concatenated and analyzed with FCS Express 7 (DeNovo Software, Pasadena, CA) as described previously.^26,30^

### Statistical Analysis

All analyses were performed using Prism 10 (GraphPad, La Jolla, CA). The data were tested for normality using the Shapiro-Wilk test. Statistical significance for differences was determined using either a parametric unpaired t-test or a one-way ANOVA followed by Tukey’s Multiple comparisons test, as appropriate. OCR data were analyzed using a two-way repeated measures ANOVA. All data are presented as mean±SEM, and differences were considered significant at p < 0.05.

## RESULTS

### NOX4-Dependent Mitochondrial Oxidative Stress Is Increased in Human Abdominal Aortic Aneurysm (AAA)

Increased NOX4 expression was implicated in the development of AAA in humans.^31^ To investigate the relationship between NOX4-derived ROS generation and AAA development, we analyzed NOX4 expression in the aortic tissues of human control subjects and AAA patients. Western blot analysis of aortic lysates revealed an 11-fold increase in NOX4 protein levels in AAA samples compared to control aortas (Figures 1A & B). In agreement with this finding, immunofluorescence analysis showed a more than 35-fold increase in NOX4 immunoreactivity in cryosections from AAA tissue (Figures 1C & D). Furthermore, colocalization analysis indicated a 14-fold increase in NOX4 signal overlapping with the mitochondrial marker ATP5G in AAA sections, suggesting enhanced mitochondrial expression of NOX4 (Figures 1C & E). Additionally, mitochondrial ROS levels, assessed by MitoSOX fluorescence, were markedly elevated in AAA tissue compared to control samples (Figures 1F & G).

**Figure 1.**
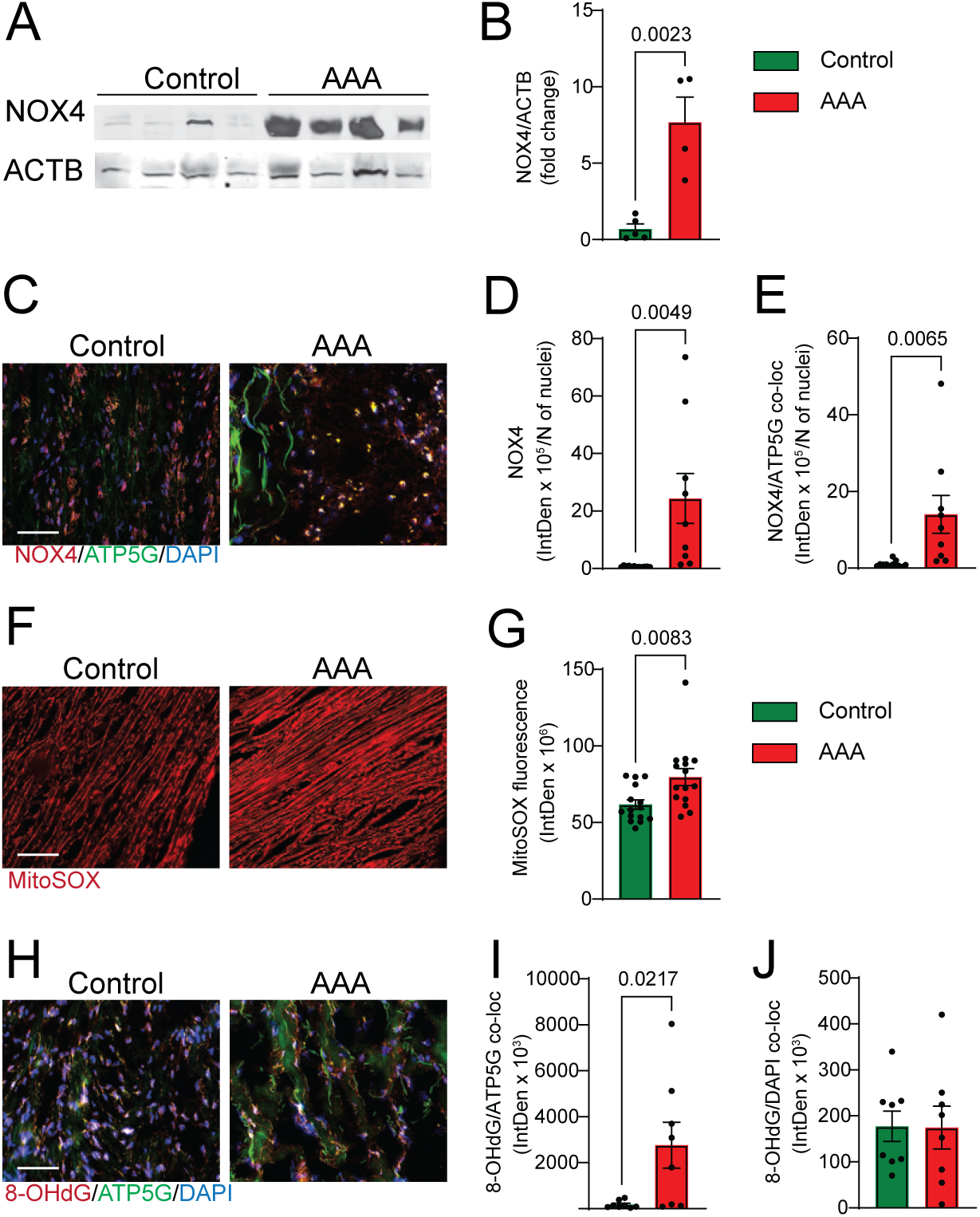
Increased expression of NOX4 in human abdominal aortic aneurysm (AAA) samples is associated with markers of mitochondrial oxidative stress. *A*: Western blot analysis of NOX4 protein expression in lysates from human control aorta and AAA samples. *B*: Quantification of NOX4 protein expression in lysates from human control aorta and AAA samples, presented as fluorescence intensity fold change over control, adjusted for ACTB levels (mean ± SEM, n = 4). *C*: Representative fluorescence microscopy images of frozen sections of human control aorta and AAA samples, stained for immunoreactive NOX4 (red) and ATP5G (green), counterstained with DAPI (blue). The scale is 100 µm. *D*: Quantification of immunoreactive NOX4 expression in control and AAA samples, represented as fluorescence integrated density adjusted for the number of stained nuclei (mean ± SEM, n = 9). *E*: Quantification of fluorescence colocalization of NOX4 and the mitochondrial protein ATP5G in control and AAA samples, presented as fluorescence integrated density adjusted for the number of stained nuclei (mean ± SEM, n = 9). *F*: Representative fluorescence microscopy images of MitoSOX staining in frozen human control and AAA aortic sections. Scale bar: 100 µm. *G*: Quantification of MitoSOX fluorescence in control and AAA aortic samples, represented as fluorescence integrated density (mean ± SEM, n =15). *H*: Representative fluorescence microscopy images of frozen sections of human control and AAA aortas stained for immunoreactive 8-OHdG (red) and ATP5G (green), counterstained with DAPI (blue). Scale bar: 100 µm. *I*: Quantification of fluorescence colocalization of 8-OHdG and DAPI in control and AAA samples, presented as fluorescence integrated density (mean ± SEM, n = 8). *J*: Quantification of fluorescence colocalization of 8-OHdG and ATP5G in control and AAA aortas, presented as fluorescence integrated density (mean ± SEM, n = 8).

Increased levels of NOX4-derived ROS are linked to oxidative damage to both nuclear and mitochondrial DNA.^16,26^ Therefore, we examined DNA oxidation in the human aorta cryosections. Immunofluorescence staining for 8-hydroxy-2’-deoxyguanosine (8-OHdG), a well-known marker of oxidative DNA damage, revealed a significantly higher colocalization of 8-OHdG with ATP5G in aortic samples from patients with AAA, suggesting enhanced mitochondrial DNA damage (Figures 1H & I). In contrast, we did not observe a significant difference in the colocalization of 8-OHdG with nuclear DAPI staining between AAA and control aortic tissues (Figures 1H & J), suggesting that oxidative DNA damage was predominantly localized to mitochondria in AAA. These data demonstrate that mitochondrial NOX4 expression is significantly upregulated in human AAA, contributing to increased mitochondrial ROS and DNA damage, which suggests a key role in the disease pathogenesis.

Increased NOX4 Expression Promotes AAA Development in *Apoe^-/-^* Mice

Previous studies have demonstrated elevated NOX4 expression in atherosclerotic arteries and its association with adverse disease progression in humans and murine models.^16,32^ To investigate the role of NOX4-dependent mitochondrial oxidative stress in AAA pathogenesis, we assessed aneurysm formation in Ang II-infused *Apoe^-/-^*, *Nox4*TG/*Apoe*^-/-^, and *Nox4^-/-^/Apoe^-/-^* mice over 28 days. Gross examination of the aortas revealed AAA development in 44% of *Apoe^-/-^* mice and 83% of *Nox4*TG/*Apoe*^-/-^ mice, whereas no aneurysms were observed in *Nox4^-/-^/Apoe^-/-^* mice (Figures 2A & B).

**Figure 2.**
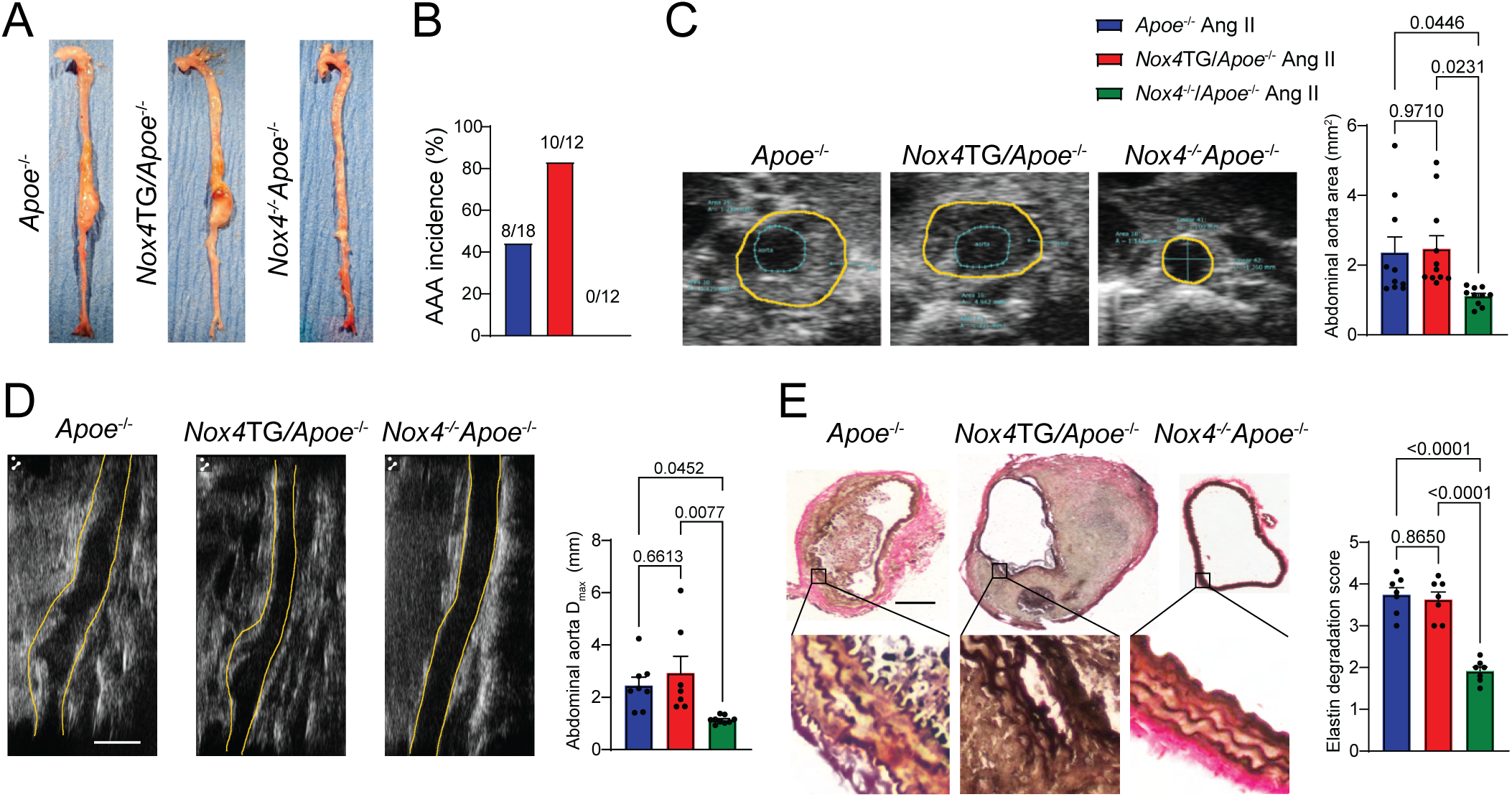
NOX4 levels correlate with the incidence of AAA in *Apoe*^-/-^, *Nox4*TG/*Apoe*^-/-^, and *Nox4*^-/-^/*Apoe*^-/-^ mice treated with Ang II for 28 days. *A*: Representative images of gross aorta from mice treated with Ang II for 28 days. *B*: The incidence of AAA in the treated mice. *C*: Representative transverse ultrasound images and quantification of abdominal aorta area in mice treated with Ang II for 28 days. The scale is 1 mm. Data presented are mean ± SEM, n = 10. *D*: Representative sagittal ultrasound images and quantification of the maximum diameter of the abdominal aorta in mice treated with Ang II for 28 days. The scale is 2 mm. Data are mean ± SEM, n = 10. *E*: Representative microscopy images and quantification of Verhoeff-van Gieson elastic stain in transverse sections of the abdominal aorta from mice treated with Ang II for 28 days. Higher magnification insets (black rectangles) are shown in the lower panel. The scale is 100 µm. Data represent the elastin degradation score (mean ± SEM, n = 7).

Ultrasound imaging performed on day 28 post-Ang II infusion showed significantly enlarged abdominal aorta area in *Apoe^-/-^* and *Nox4*TG/*Apoe*^-/-^ mice compared to *Nox4^-/-^ /Apoe^-/-^* mice (Figure 2C). Similarly, the maximal abdominal aorta diameter was significantly increased in *Apoe^-/-^* and *Nox4*TG/*Apoe*^-/-^ mice relative to *Nox4^-/-^/Apoe^-/-^* mice (Figure 2D). Histological analysis using Verhoeff–Van Gieson (VVG) staining of abdominal aortic cryosections revealed elastin degradation and disrupted laminae integrity in *Apoe^-/-^* and *Nox4*TG/*Apoe*^-/-^ mice. In contrast, the elastin structure remained largely intact in *Nox4^-/-^/Apoe^-/-^* mice (Figure 2E). Quantification of elastin degradation scores confirmed significantly greater elastolysis in *Apoe^-/-^* and *Nox4*TG/*Apoe*^-/-^ mice compared to *Nox4^-/-^/Apoe^-/-^* controls (Figure 2E).

### Increased Mitochondrial Oxidative Stress Drives Phenotypic Modulation of Aortic Wall Cells in *Apoe^⁻/⁻^* Mice with AAA

Elevated NOX4 expression in the vascular wall under pathological conditions enhances mitochondrial ROS production, promoting detrimental cellular phenotypic transitions. ^16,21,26^ To investigate the relationship between mitochondrial oxidative stress, vascular cell phenotype changes, and AAA development, we analyzed ROS levels in abdominal aortic cryosections from *Apoe^-/-^*, *Nox4*TG/*Apoe*^-/-^ and *Nox4^-/-^/Apoe^-/-^* mice following 28 days of Ang II infusion. DHE fluorescence staining revealed significantly elevated intracellular ROS levels in *Nox4*TG/*Apoe*^-/-^ compared to *Nox4^-/-^/Apoe^-/-^* mice aortas (Figure 3A). Similarly, mitochondrial ROS levels, assessed using MitoSOX fluorescence, were significantly increased in aortic cells from Ang II-treated *Apoe^-/-^* and *Nox4*TG/*Apoe*^-/-^ mice compared with *Nox4^-/-^/Apoe^-/-^* mice (Figure 3B). In line with these findings, immunofluorescence analysis showed significantly higher levels of immunoreactive 8-OHdG co-localized with the mitochondrial protein TOMM20 in *Apoe^-/-^* and *Nox4*TG/*Apoe*^-/-^ compared with *Nox4^-/-^/Apoe^-/-^* mice aortas (Figure 3C).

**Figure 3.**
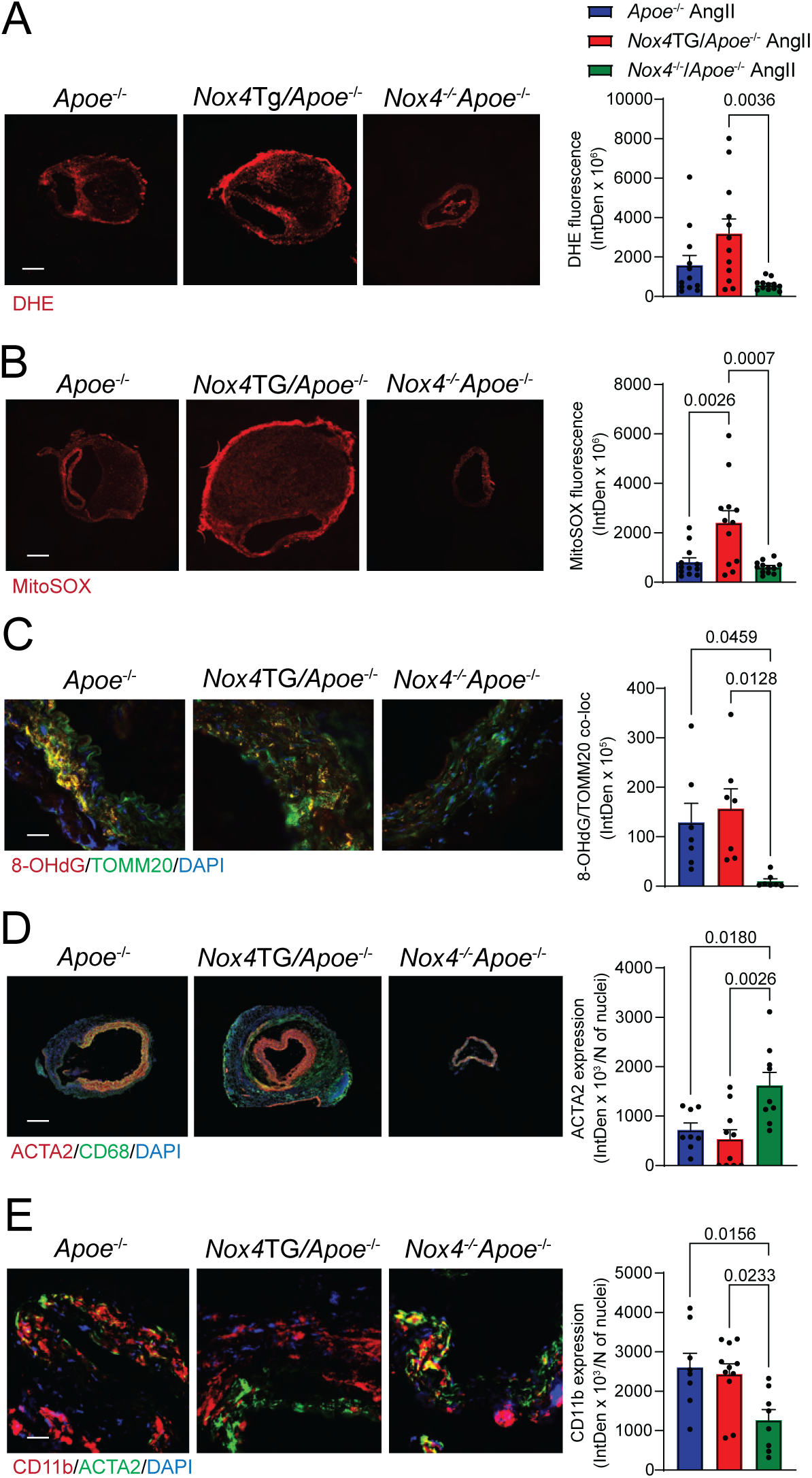
NOX4-derived oxidative stress is associated with phenotypic changes in aortic wall cells in *Apoe*^-/-^, *Nox4*TG/*Apoe*^-/-^, and *Nox4*^-/-^/*Apoe*^-/-^ mice treated with Ang II for 28 days. *A & B*: Representative fluorescence microscopy images and quantification of DHE (*A*) and MitoSOX (*B*) fluorescence in transverse abdominal aorta sections from mice treated with Ang II for 28 days. The scale bar represents 100 µm. Data are presented as fluorescence integrated density (mean ± SEM, n = 12). *C*: Representative fluorescence microscopy images and quantification of the colocalization of 8-OHdG (red) and TOMM20 (green), with DAPI (blue) used for counterstaining. The scale bar is 10 µm. Data are fluorescence integrated density (mean ± SEM, n = 7). *D*: Representative fluorescence microscopy images and quantification of ACTA2 expression in transverse abdominal aorta sections from mice treated with Ang II for 28 days. These sections were stained for ACTA2 (red) and CD68 (green) and counterstained with DAPI (blue). The scale bar is 100 µm, and data are expressed as fluorescence integrated density (mean ± SEM, n = 9). *E*: Representative fluorescence microscopy images and quantification of CD11b expression in transverse abdominal aorta sections from mice treated with Ang II for 28 days. The sections were stained for CD11b (red), ACTA2 (green), and counterstained DAPI (blue). The scale is 100 µm. Data are presented as fluorescence integrated density (mean ± SEM, n = 9).

Increased vascular oxidative stress in hypercholesterolemic mice has been associated with the phenotypic transformation of smooth muscle cells into macrophage-like cells.^29^ In line with this, immunofluorescence staining revealed a significant reduction in the expression of the SMC contractile marker, α-smooth muscle actin (ACTA2), in abdominal aortic sections from Ang II–treated *Apoe^-/-^* and *Nox4*TG/*Apoe*^-/-^ mice compared to *Nox4^-/-^/Apoe^-/-^* mice (Figure 3D). The inflammatory leukocyte marker CD11b expression significantly increased in the aortic sections of *Apoe^-/-^* and *Nox4*TG/*Apoe*^-/-^ (Figure 3E). These results suggest that NOX4-driven ROS production and oxidative damage are linked to the loss of the contractile SMC phenotype and the development of inflammatory characteristics, potentially contributing to AAA pathogenesis.

NOX4-Dependent Mitochondrial Dysfunction Mediates Ang II–Induced Pro-Inflammatory Phenotype Switching in Aortic SMCs

To investigate the role of NOX4 in mitochondrial dysfunction, we measured mitochondrial ROS production in primary aortic VSMCs isolated from wild-type, *Nox4*TG, and *Nox4^-/-^* mice treated with either vehicle or Ang II. Under basal conditions, MitoSOX fluorescence—a marker of mitochondrial ROS—did not differ significantly among genotypes (Figure 4A). However, Ang II stimulation led to a marked increase in mitochondrial ROS levels in all cell types, with the highest fluorescence observed in *Nox4*TG VSMCs, the lowest in *Nox4^-/-^* cells, and intermediate levels in wild-type cells (Figure 4A). In line with this, ELISA-based quantification of 8-OHdG revealed a significant increase in oxidative DNA damage levels in Ang II–treated *Nox4*TG VSMCs compared to wild-type and *Nox4^-/-^*cells (Figure 4B).

**Figure 4.**
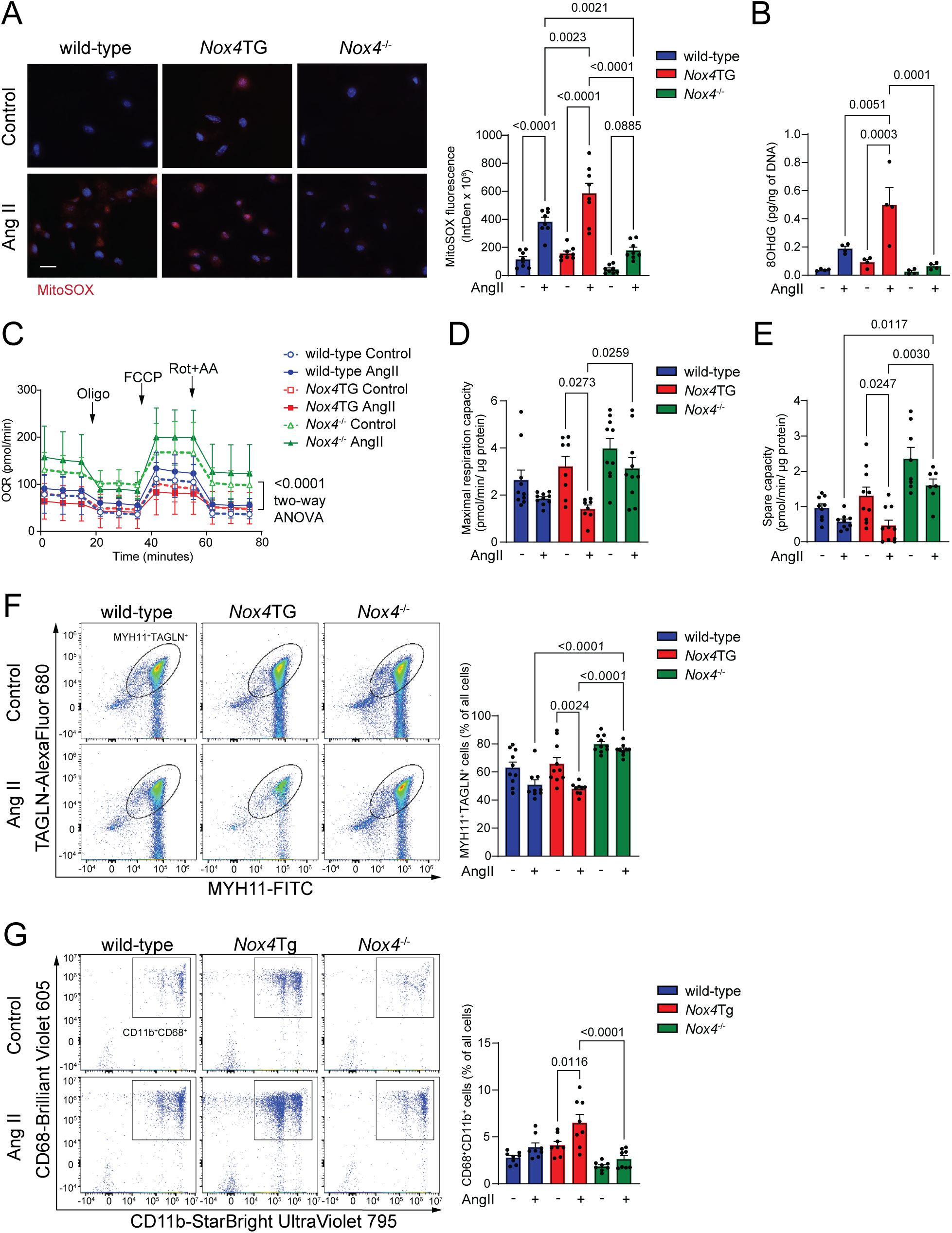
Higher levels of NOX4 expression are associated with mitochondrial oxidative stress, DNA damage, and dysfunction, leading to phenotypic change in VSMC treated with Ang II from wild-type, *Nox4*TG, and *Nox4*^-/-^ mice. *A*: Representative fluorescence microscopy images and quantification of MitoSOX fluorescence in VSMCs treated with either a vehicle or 100 µM Ang II for 30 min. The scale bar is 2 µm. Data represent the integrated density of MitoSOX fluorescence (mean ± SEM, n = 8). *B*: The amount of 8-OHdG was measured by ELISA in total DNA from VSMCs treated with a vehicle or 100 µM Ang II for 24 h (mean ± SEM, n = 4). *C*: The oxygen consumption rate (OCR) was measured in VSMCs treated with a vehicle or Ang II for 24 h using an Agilent Seahorse XF96 analyzer (mean ± SEM, n = 12). *D*: Mitochondrial maximal respiration capacity was derived from OCR measurements in VSMCs treated with either a vehicle or Ang II for 24 h (mean ± SEM, n = 10). *E*: Mitochondrial spare capacity was also derived from OCR measurements in VSMCs treated with either a vehicle or Ang II for 24 h (mean ± SEM, n= 10). *F*: Flow cytometry analysis and quantification of MYH11^+^TAGLN^+^ cells in VSMCs treated with either a vehicle or Ang II for 24 h (mean±SEM, n=10). *G*: Flow cytometry analysis and quantification of CD11b^+^CD68^+^ cells in VSMCs treated with either a vehicle or Ang II for 24 h (mean ± SEM, n = 10).

Since mitochondrial oxidative stress is strongly linked to impaired mitochondrial respiration, ^26,33^ we assessed mitochondrial function by measuring the oxygen consumption rate (OCR) using the Seahorse XF analyzer. Treatment with Ang II reduced OCR across all cell types. Interestingly, *Nox4^-/-^* VSMCs exhibited the highest OCR, while *Nox4*TG VSMCs displayed the lowest rates at baseline and after Ang II exposure (Figure 4C). Similarly, both maximal respiration and spare respiratory capacity decreased in response to Ang II in all groups, but they remained significantly higher in *Nox4^-/-^* VSMCs compared to wild-type and *Nox4*TG cells (Figures 4D & E). These results indicate that NOX4 expression levels directly impact mitochondrial oxidative stress and functional capacity in VSMCs.

To investigate whether NOX4-driven mitochondrial dysfunction contributes to the phenotypic modulation of SMCs, we analyzed changes in marker expression in Ang II- treated VSMCs using flow cytometry. Ang II exposure significantly decreased the proportion of cells expressing contractile SMC markers MYH11 and TAGLN in wild-type and *Nox4*TG VSMCs (p < 0.01 for *Nox4*TG; Figure 4F). In contrast, MYH11^+^TAGLN^+^ cell populations remained stable in *Nox4^-/-^* VSMCs and were significantly higher than in the other groups post-Ang II treatment. Additionally, the percentage of cells expressing macrophage-specific markers CD68 and CD11b significantly increased in Ang II-treated *Nox4*TG VSMCs, significantly exceeding levels observed in similarly treated *Nox4^-/-^* cells (Figure 4G). These findings indicate that NOX4-dependent mitochondrial ROS generation and dysfunction drive a pro-inflammatory phenotypic shift in VSMCs, characterized by downregulation of contractile SMC markers and upregulation of macrophage markers in response to Ang II.

### Reciprocal Upregulation of DNase II Expression in AAA Correlates with Increased NOX4 Expression and Mitochondrial DNA Damage

Mitochondrial DNA oxidative damage and subsequent release into the cytosol can activate innate immune responses and upregulate deoxyribonuclease II (DNase II) expression.^34^ In line with the increased mtDNA damage observed in abdominal aortas of Ang II-treated *Nox4*TG/*Apoe^-/-^* mice, DNase II protein expression was significantly elevated in ACTA2^+^ regions of AAA aortic sections from both *Apoe^-/-^* and *Nox4*TG/*Apoe^-/-^* mice compared with *Nox4^-/-^/Apoe^-/-^*controls (1.66- and 2.17-fold increases, respectively; Figure 5A).

**Figure 5.**
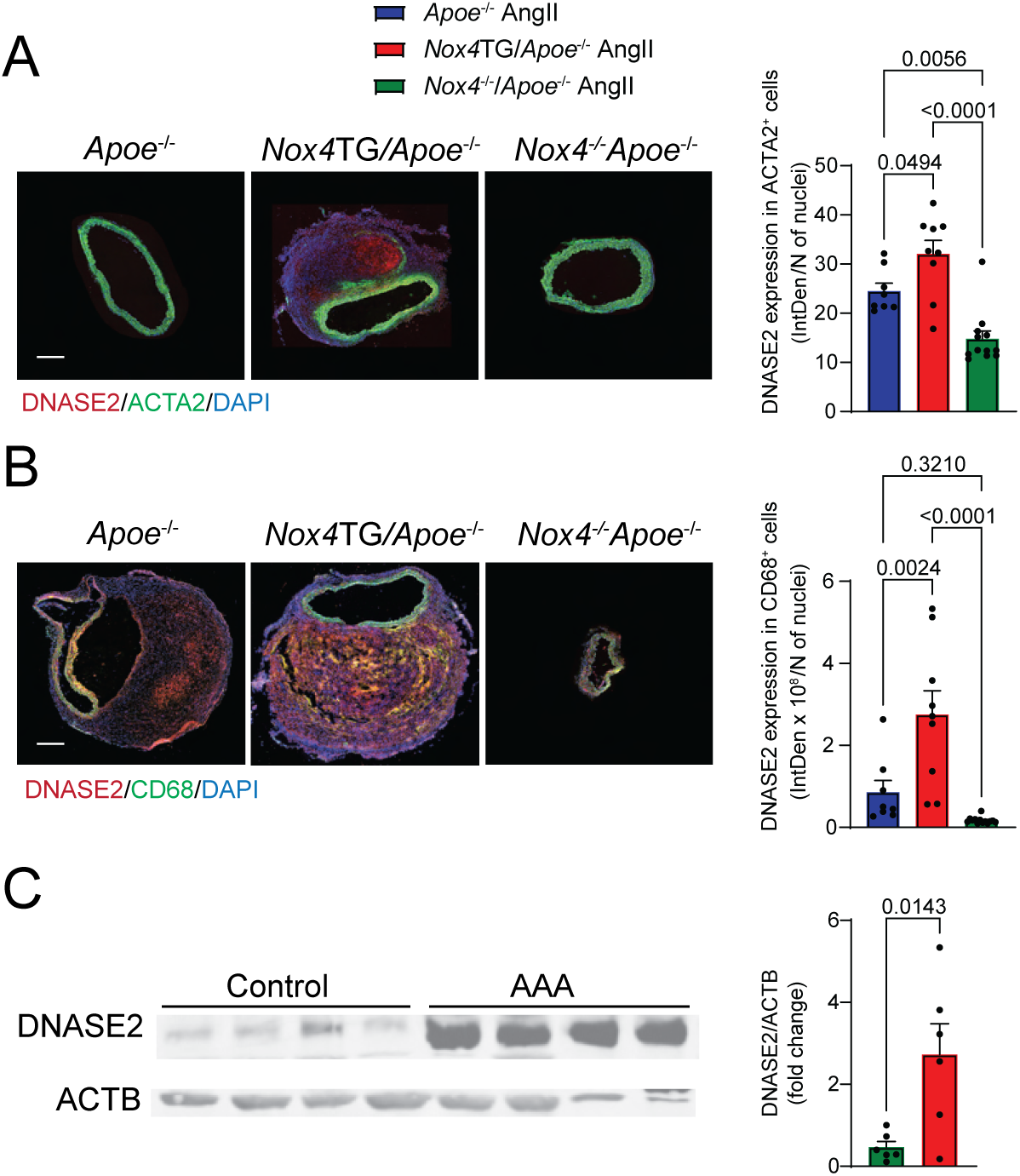
A reciprocal increase in DNASE2 expression levels is associated with elevated NOX4 levels in mice and human AAA. *A* & *B*: Representative fluorescence microscopy images and the quantification of fluorescence colocalization of DNASE2 (red) with ACTA2 (green) (*A*) and with CD68 (green) (*B*) These images were obtained from transverse abdominal aorta sections of *Apoe*^-/-^, *Nox4*TG/*Apoe*^-/-^, and *Nox4*^-/-^/*Apoe*^-/-^ mice treated with Ang II for 28 days. The scale bar is 100 µm, and the data are expressed as fluorescence integrated density (mean ± SEM, n = 9). *C*: Western blot analysis of DNASE2 expression in the protein lysates from human aorta samples, comparing control and AAA samples. The data are expressed as fluorescence intensity fold change over control, adjusted for ACTB levels (mean ± SEM, n = 6).

Furthermore, DNase II expression within CD68^+^ macrophage-enriched areas was highest in *Nox4*TG/*Apoe^-/-^* aneurysms, intermediate in *Apoe^-/-^* aortas, and lowest in *Nox4^-/-^*/*Apoe^-/-^* aortas (16.32- and 5.08-fold increases, respectively; Figure 5B). Consistent with these murine data, DNase II protein levels were significantly elevated in lysates from human AAA samples compared to control aortic tissue (Figure 5C).

These findings suggest that NOX4-mediated mtDNA damage and cytosolic leakage may drive DNase II upregulation, contributing to the inflammatory milieu and AAA pathogenesis.

### Mitochondrial DNA Damage Activates Cytosolic DNA Sensing in Ang II-treated *Nox4*TG VSMC and *Nox4*TG/*Apoe*^-/-^ Mouse Aortas

Previous studies have demonstrated that nuclear and mitochondrial DNA damage, as well as cytosolic leakage, trigger the activation of the DNA-sensing STING pathway, which is crucial for SMC phenotypic changes and the development of AAA.^35^ The activation of the cytosolic DNA-sensing cGAS-STING pathway also plays a role in pro- inflammatory responses in atherosclerotic aortas.^36,37^ To investigate the role of NOX4- dependent oxidative DNA damage in activating the cytosolic DNA-sensing pathway and its impact on AAA development, we first examined cGAS and STING expression in Ang II-treated cultured VSMCs isolated from wild-type, *Nox4*TG, and *Nox4*^-/-^ mice.

Western blot analysis revealed a significant increase in cGAS protein levels in *Nox4*TG VSMC, while no such increase was observed in wild-type or *Nox4^-/-^* cells (Figures 6A & B). The cytoplasmic levels of cGAMP, a product of cGAS activation, were significantly elevated in Ang II-treated *Nox4*TG compared to *Nox4^-/-^* cells. However, they were not substantially different from those in wild-type cells (Figure 6C). This activation was accompanied by significantly higher STING protein levels in the Ang II-treated transgenic VSMC lysates than in knockout cells (Figures 6A & D).

**Figure 6.**
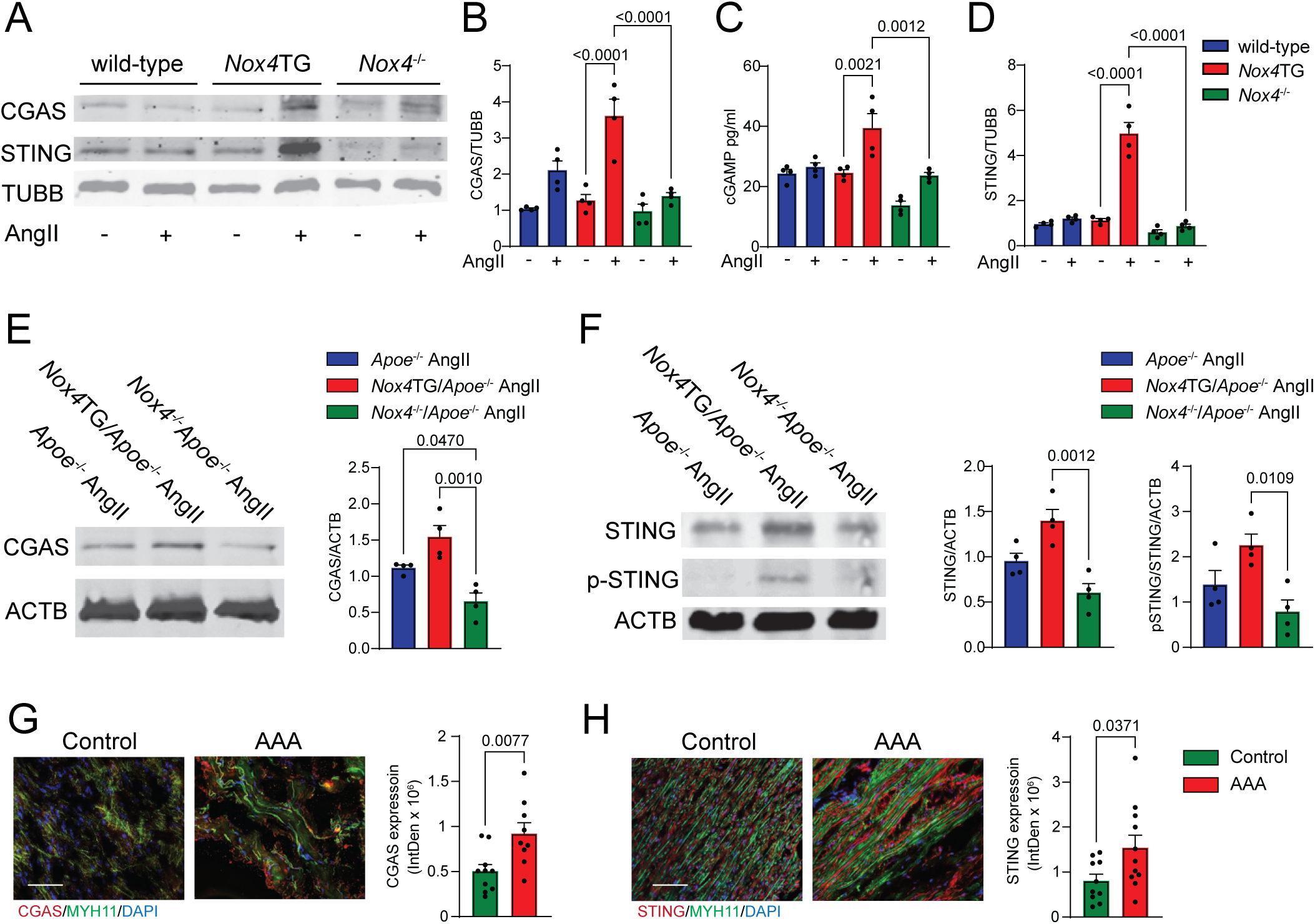
Mitochondrial DNA damage activates the cytosolic DNA sensing pathway in Ang II-treated *Nox4*TG VSMC, *Nox4*TG/*Apoe*^-/-^ mice abdominal aortas, and human AAA samples. *A*: Western blot analysis of CGAS and STING expression in protein lysates from vehicle or Ang II-treated VSMCs isolated from wild-type, *Nox4*TG, and *Nox4*^-/-^ mice. *B*: The quantification of CGAS protein expression in vehicle or Ang II-treated VSMCs. The data is fluorescence intensity fold change over vehicle- treated wild-type cells, adjusted for TUBB levels (mean ± SEM, n = 4). *C*: CGAMP levels were measured using ELISA in cell lysates from vehicle or Ang II-treated VSMCs (mean ± SEM, n = 4). *D*: The quantification of STING protein expression was analyzed with Western blot in vehicle or Ang II-treated VSMCs. Data are presented as fluorescence intensity fold change over vehicle-treated wild-type cells, adjusted for TUBB levels (mean ± SEM, n=4). *E*: Western blot analysis and quantification of CGAS protein levels in the abdominal aorta protein lysates from *Apoe*^-/-^, *Nox4*TG/*Apoe*^-/-^, and *Nox4*^-/-^/*Apoe*^-/-^ mice treated with Ang II. STING protein expression was quantified with Western blot in vehicle or Ang II-treated VSMCs. Data are presented as fluorescence intensity fold change over *Apoe*^-/-^ mice, adjusted for ACTB levels (mean ± SEM, n = 4). *F*: Western blot analysis and quantification of STING and phospho-STING levels were performed on abdominal aorta protein lysates from *Apoe*^-/-^, *Nox4*TG/*Apoe*^-/-^, and *Nox4*^-/-^/*Apoe*^-/-^ mice treated with Ang II. Data are presented as fluorescence intensity fold change over *Apoe*^-/-^ mice, adjusted for ACTB levels (mean ± SEM, n = 4). *G & H*: Representative fluorescence microscopy images and quantification of CGAS (*G*) and STING (*H*) expression in human control aorta and AAA frozen sections. These sections were stained for immunoreactive CGAS (*G*) or STING (*H*) (red) and MYH11 (green), with DAPI (blue) used for counterstaining. The scale bar is 100 µm. Data are presented as fluorescence integrated density (mean ± SEM, n = 9).

Similarly, Western blot analysis of abdominal aortic tissue lysates from mice treated with Ang II for 28 days showed significantly higher cGAS levels in *Apoe*^-/-^ and *Nox4*TG/*Apoe*^-/-^ aortas compared to *Nox4*^-/-^/*Apoe*^-/-^ tissue (Figure 6E). In line with observation in cultured cells, STING and its activated form, phospho-STING, were expressed at the highest levels in abdominal aorta tissue from *Nox4*TG/*Apoe*^-/-^ compared to *Nox4*^-/-^/*Apoe*^-/-^ mice (Figure 6F).

These results suggest that NOX4-dependent mitochondrial oxidative stress and DNA damage activate the cytosolic DNA-sensing pathway, contributing to aneurysm development and progression. Supporting this, the levels of immunoreactive cGAS and STING proteins were significantly higher in the human AAA samples compared to control aorta tissue (Figures 6G &H), underscoring the involvement of this pathway in aneurysm pathophysiology.

### Inflammation-driven Alterations in Vascular Cell Phenotype Characterize AAA

Activation of the cGAS-STING pathway promotes an inflammatory phenotype in mouse vascular cells.^37,38^ To determine whether similar changes occur in human AAAs, we analyzed the cellular composition and inflammatory cytokine expression in control and AAA tissue samples. Immunofluorescence microscopy revealed a significant increase in the inflammatory macrophage marker CD68 and a corresponding decrease in the SMC contractile marker ACTA2 in AAA tissues compared to controls (3.2-fold increase in CD68 and 4.6-fold reduction in ACTA2; Figure 7A). Consistent with these findings, levels of the pro-inflammatory cytokines IL-1β and IL-6 were significantly elevated in AAA samples relative to controls (3.0-fold and 3.2-fold increases, respectively; Figures 7B & 7C). These results support the hypothesis that a phenotypic switch of VSMCs toward a pro-inflammatory state contributes to AAA pathogenesis.

**Figure 7.**
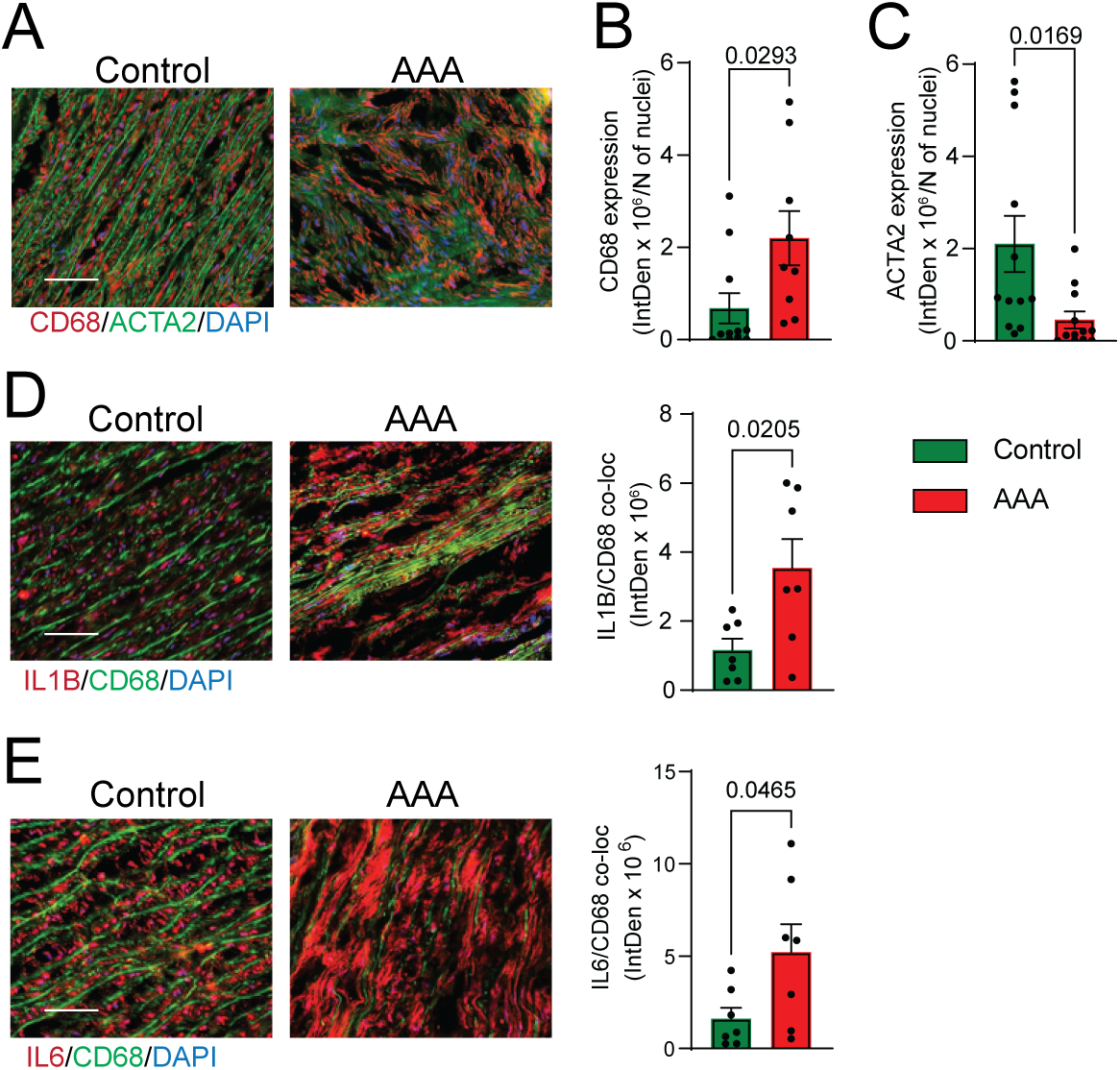
AAA is associated with increased expression of macrophage and inflammatory markers in the human abdominal aorta. *A*: Representative fluorescence microscopy images of human control aorta and AAA frozen sections, stained for immunoreactive CD68 (red) and ACTA2 (green), with DAPI used as a counterstain (blue). The scale bar is 100 µm. *B & C*: Quantification of immunoreactive CD68 (*B*) and ACTA2 (*C*) expression in control and AAA samples. The data are presented as fluorescence integrated density, adjusted for the number of stained nuclei ((mean ± SEM, n = 9). *D & E*: Representative fluorescence microscopy images and quantification of colocalization of IL1b (*D*) or IL6 (*E*) (red) with CD68 (green) expression in human control aorta and AAA frozen sections. The scale bar is 100 µm. The data are fluorescence integrated density (mean ± SEM, n = 7).

### Increased Mitochondrial NOX4 Expression is Associated with Pro-Inflammatory Macrophage-like Phenotype in Aortic SMC During AAA Development

To determine the role of NOX4-dependent in activating interferon response pathways and inducing a pro-inflammatory phenotype in SMCs during the development of AAA, we conducted spectral flow cytometry analysis of aortic tissue from Ang II-treated *Apoe*^-/-^, *Nox4*TG/*Apoe*^-/-^, and *Nox4*^-/-^/*Apoe*^-/-^ mice after 28 days, using antibodies against contractile and inflammatory proteins.

We observed a reduction in the proportion of MYH11^+^ cells in the aortas of *Apoe*^-/-^ and *Nox4*TG/*Apoe*^-/-^ mice in comparison to *Nox4*^-/-^/*Apoe*^-/-^ mice (84.2% and 84% vs. 90.1%, respectively; Figure 8A). Conversely, the proportion of CD45^+^CD68^+^CD11b^+^ inflammatory cells was significantly higher in *Apoe*^-/-^ mice with elevated NOX4 expression compared to the *Nox4^-/-^/ Apoe*^-/-^ mice (14% and 16.5% vs. 8.7%, respectively; Figure 8B). Notably, the fraction of MYH^+^ cells co-expressing the inflammatory markers CD68 and CD11b was substantially higher in *Apoe*^-/-^ and *Nox4*TG/*Apoe*^-/-^ mice than in *Nox4*^-/-^/*Apoe*^-/-^ mice (11.5% and 11.1% vs. 5.3%, respectively; Figure 8C). These findings indicate that NOX4 promotes a phenotypic transition of SMCs towards a pro-inflammatory, macrophage-like state, contributing to the pathogenesis of AAA.

**Figure 8.**
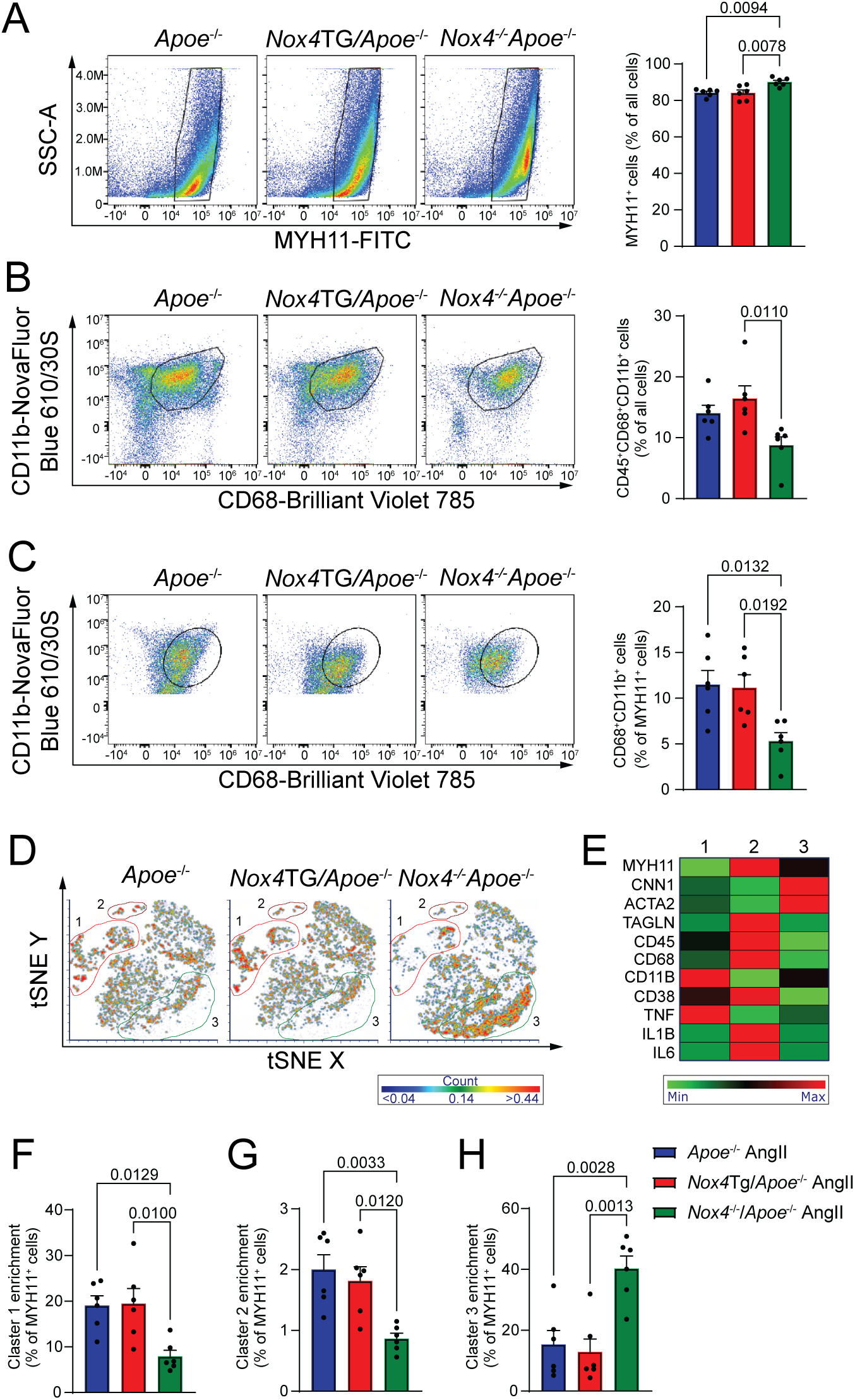
Increased expression of macrophage markers in SMC within the abdominal aorta of *Apoe*^-/-^, *Nox4*TG/*Apoe*^-/-^, and *Nox4*^-/-^/*Apoe*^-/-^ mice treated with Ang II. *A & B*: Flow cytometry analysis of single-cell suspension from the abdominal aorta of mice treated with Ang II. Panel *A* presents the quantification of MYH11^+^ cells, while Panel *B* shows the quantification of CD45^+^CD68^+^CD11b^+^ cells. The data represent the cell fraction of all aortic cells (mean ± SEM, n=6). *C*: Flow cytometry analysis and quantification of abdominal aorta single-cell suspension from Ang II-treated mice showing the proportion of MYH11^+^ cells expressing macrophage markers. This data shows the fraction of CD68^+^CD11b^+^ cells as a percentage of MYH11^+^ cells (mean ± SEM, n = 6). *D*: A t-SNE clustering analysis of flow cytometry data from the abdominal aorta of Ang II-treated mice was performed. MYH11^+^ cells were clustered based on the expression of CNN1, ACTA2, TAGLN, CD45, CD68, CD11b, CD38, TNFα, IL1b, and IL6, resulting in distinct clusters presented on the t-SNE plot. *E*: A heat map representation illustrates the relative expression of SMC, macrophage, and inflammatory markers based on the mean fluorescence intensity for each cluster. *F* & *H*: The relative abundance of distinct clusters as a proportion of MYH11^+^ cells (mean ± SEM, n = 6).

To identify subsets of VSMCs with distinct contractile or macrophage-like phenotypes, we performed unbiased multidimensional t-distributed stochastic neighbor embedding (t-SNE) analysis of MYH11⁺ abdominal aorta cells flow cytometry (Figure 8D). We identified three distinct clusters whose relative abundance varied significantly among *Apoe*^-/-^, *Nox4*TG/*Apoe*^-/-^, and *Nox4*^-/-^/*Apoe*^-/-^ aortas. Clustering was based on expression of canonical SMC contractile markers (MYH11, CNN1, ACTA2, and TAGLN), macrophage markers (CD45, CD68, CD11b, and CD38), and inflammatory cytokines (TNF, IL-1β, and IL-6) (Figures 8D & 8E).

Cluster 1 comprises macrophage-like SMCs with elevated CD11b and TNF expression. This cluster was significantly enriched in *Apoe*^-/-^ and *Nox4*TG/*Apoe^-/-^* compared to *Nox4*^-/-^/*Apoe*^-/-^ mouse aortas (19.1% and 19.5% vs. 7.9% of MYH11^+^ cells, respectively; Figures 8E & 8F). Cluster 2, representing a smaller population of macrophage-like SMCs with higher levels of CD45, CD68, IL-1β, and IL-6, was also significantly more abundant in *Apoe*^-/-^ and *Nox4*TG/*Apoe^-/-^* than in *Nox4*^-/-^/*Apoe*^-/-^ aortas (2.0% and 1.8% vs. 0.86%, respectively; Figures 8E & 8G). In contrast, Cluster 3, composed of contractile SMCs with high expression of CNN1 and ACTA2, was significantly enriched in *Nox4*^-/-^/*Apoe*^-/-^ aortas relative to *Apoe*^-/-^ and *Nox4*TG/*Apoe***^⁻/⁻^** tissues (40.3% vs. 15.3% and 12.9%, respectively; Figures 8E & 8H). These findings indicate that NOX4 is pivotal in promoting the phenotypic transition of SMCs toward a pro-inflammatory, macrophage-like state during AAA development. Conversely, genetic deletion of *Nox4* preserves the contractile SMC phenotype and attenuates vascular inflammation, thereby protecting against AAA formation.

## DISCUSSION

The present study uncovers a pivotal role for mitochondrial NOX4-derived ROS in driving SMC phenotypic switching, inflammation, and aortic wall remodeling during AAA pathogenesis. Using genetic models, primary cell culture, and human tissue analysis, we delineate a mechanistic cascade whereby mitochondrial NOX4 promotes oxidative stress and mtDNA damage, ultimately activating redox-sensitive innate immune signaling pathways that govern SMC fate decisions and vascular homeostasis.

While NOX4 has previously been implicated in vascular pathologies, including hypertension and atherosclerosis, ^16,26,39^ our findings uniquely position mitochondrial NOX4 as a spatially and functionally distinct source of oxidative stress that orchestrates maladaptive vascular remodeling in AAA. We observed increased expression of mitochondrial NOX4 in aneurysmal tissue, which was strongly associated with elevated mitochondrial ROS, DNA oxidation, and dysfunction. Our data are supported by the observation that AAA-derived VSMCs exhibit persistent mitochondrial dysfunction characterized by increased fission, elevated ROS production, impaired antioxidant response to Ang II, and accumulation of oxidative DNA damage.^40^

The oxidative milieu in mitochondria triggers the leakage of mtDNA into the cytosol, a critical upstream trigger of the cGAS-STING signaling axis.^35^ Activation of this DNA- sensing pathway induces pro-inflammatory and interferon responses. It enhances the activity of IRF3 (Interferon Regulatory Factor 3), which recruits the histone methyltransferase EZH2 (enhancer of zeste homolog 2) to the promoters of contractile genes. This process results in the addition of the repressive H3K27me3 epigenetic mark, potentially silencing contractile genes and driving the transition SMCs into a pro- inflammatory, macrophage-like phenotype.^38^

Our data show a significant increase in the inflammatory macrophage marker CD68 and a notable decrease in the SMC contractile marker ACTA2, along with elevated levels of pro-inflammatory cytokines IL-1β and IL-6 in human AAA samples when compared to controls. Importantly, the activation of the NLRP3 inflammasome in macrophages, through a mechanism-dependent mtROS, induces the release of IL-1β and the upregulation of matrix metalloproteinases, resulting in vascular wall remodeling and the formation of AAA.^41^

Consistent with the elevated IL-6 levels we observed in human AAA tissues, recent Mendelian randomization analyses provide robust genetic evidence that increased IL-6 signaling causally contributes to AAA development.^42^ This study supports the inhibition of the IL-6 pathway as a promising therapeutic strategy specific to AAA pathophysiology, distinct from other aneurysmal diseases. Disrupting the cGAS-STING signaling pathway helped maintain the contractile phenotype of SMCs in vitro and protected against Ang II-induced AAA formation in *Sting^-/-^* mice, highlighting its role in redox-sensitive epigenetic reprogramming.^38^ Concurrently, we observed increased expression of DNase II in regions rich in ACTA2⁺ SMCs and CD68⁺ macrophages, suggesting compensatory activation of nucleic acid clearance pathways in response to the buildup of cytosolic mtDNA.

Our data align with and extend prior reports that highlight the phenotypic plasticity of vascular SMCs in the pathogenesis of AAA. Recent single-cell and lineage-tracing studies have shown that SMCs can adopt distinct inflammatory and phagocytic phenotypes in response to stress, contributing to immune cell recruitment, extracellular matrix degradation, and wall weakening.^38,43,44^ We provide direct evidence that NOX4- dependent mitochondrial ROS is a major driver of this phenotypic reprogramming.

Notably, we demonstrate that NOX4 overexpression in the mitochondria exacerbates AAA formation in Ang II-infused *Apoe^-/-^* mice, whereas genetic ablation of *Nox4* confers protection against aneurysm development. These findings firmly establish a causal link between mitochondrial oxidative stress and disease progression.

Our findings are further substantiated by flow cytometry and t-SNE clustering analyses, which revealed a distinct shift in the SMC population toward inflammatory subtypes in the presence of increased mitochondrial NOX4. This plasticity underscores the role of NOX4 as a molecular switch that redefines vascular cell identity under oxidative stress conditions. These results also corroborate genome-wide association studies (GWAS) that identify NOX4 as a candidate gene for AAA susceptibility,^19^ but critically, our study advances these associations by providing functional validation in both murine models and human tissues.

Considering these insights, therapeutic strategies targeting mitochondrial redox signaling warrant strong consideration. The observed maladaptive responses to NOX4 overexpression and the failure of antioxidant defenses in *Nox4*TG mice highlight the vulnerability of the aortic wall to sustained oxidative stress.^21^ Our findings mechanistically explain why *Nox4*TG mice display heightened susceptibility to Ang II- induced AAA, characterized by excessive mitochondrial ROS generation and oxidative DNA damage. These defects mirror those observed in human aneurysmal tissue and support a central role for NOX4 in disrupting mitochondrial redox homeostasis.

This mechanistic framework is strengthened by recent work by Guo et al.,^45^ who demonstrated that synthetic HDL nanoparticles restore mitochondrial function, reduce DRP1-mediated mitochondrial fission, and preserve oxidative phosphorylation in vascular SMCs, leading to a marked reduction in AAA incidence and aortic dilation in murine models. Together with our findings, this supports the notion that maintaining mitochondrial integrity in SMCs is not merely protective but essential for preventing aneurysmal degeneration. Mitochondrial dysfunction is not an epiphenomenon but a driving force in vascular remodeling and immune activation.

Additionally, our data are consistent with studies showing that the mitochondria-targeted antioxidant peptide Szeto-Schiller 31 (SS31) mitigates mtOS and ER stress, suppresses ROS generation, and reduces AAA severity in Ang II-infused *Apoe^-/-^* mice.^46^ These findings reinforce the translational potential of redox-modulating therapies, particularly those directed at mitochondria, as viable approaches for AAA management.

In conclusion, our study establishes mitochondrial NOX4 as a critical upstream mediator of oxidative damage, innate immune activation, and SMC phenotypic modulation in AAA. These insights not only elucidate the pathogenic mechanisms underpinning aneurysm formation but also highlight new opportunities for therapeutic intervention aimed at preserving mitochondrial function and redox balance. Future studies should explore the clinical relevance of these pathways in human AAA cohorts, particularly in patients with small, asymptomatic aneurysms, where pharmacological intervention could potentially alter the disease trajectory and obviate the need for surgical repair.

## Acknowledgments

We sincerely thank Dr. Marshall Runge for his invaluable support and guidance throughout this project. We also express our gratitude to Dr. Lori Isom for her generous assistance and critical review of the manuscript.

## Nonstandard Abbreviations and Acronyms

AAA: Abdominal Aortic Aneurysm
ACTA2: Alpha-Smooth Muscle Actin
AIM2: Absent in Melanoma 2
Ang II: Angiotensin II
*Apoe⁻^/^⁻*: Apolipoprotein E Knockout
cGAMP: Cyclic GMP-AMP
cGAS: Cyclic GMP-AMP Synthase
CNN1: Calponin 1
DHE: Dihydroethidium
DNASE2: Deoxyribonuclease II
ER: Endoplasmic Reticulum
FACS: Fluorescence-Activated Cell Sorting
IL-1β /IL-6: Interleukin 1 Beta / Interleukin 6
MMP2: Matrix Metalloproteinase-2
mtDNA: Mitochondrial DNA
mtOS: Mitochondrial Oxidative Stress
MYH11: Myosin Heavy Chain 11
Nox4 /*Nox4*TG /*Nox4⁻^/^⁻*: NADPH Oxidase 4/*Nox4* Transgenic/ *Nox4* Knockout
OCR: Oxygen Consumption Rate
ROS: Reactive Oxygen Species
SS31: Szeto-Schiller 31
STING: Stimulator of Interferon Genes
TAGLN: Transgelin
TOMM20: Translocase of Outer Mitochondrial Membrane 20
t-SNE: t-distributed Stochastic Neighbor Embedding
VSMC: Vascular Smooth Muscle Cell
VVG: Verhoeff–Van Gieson

